# Learning interpretable latent autoencoder representations with annotations of feature sets

**DOI:** 10.1101/2020.12.02.401182

**Authors:** Sergei Rybakov, Mohammad Lotfollahi, Fabian J. Theis, F. Alexander Wolf

## Abstract

Existing methods for learning latent representations for single-cell RNA-seq data are based on autoencoders and factor models. However, representations learned by autoencoders are hard to interpret and representations learned by factor models have limited flexibility. Here, we introduce a framework for learning interpretable autoencoders based on regularized linear decoders. It decomposes variation into interpretable components using prior knowledge in the form of annotated feature sets obtained from public databases. Through this, it provides an alternative to enrichment techniques and factor models for the task of explaining observed variation with biological knowledge. Benchmarking our model on two single-cell RNA-seq datasets, we demonstrate how our model outperforms an existing factor model regarding scalability while maintaining interpretability.

## 1. Introduction

Advances in single-cell technologies enabled constructing large cell atlases cells across different tissues and species (Regev et al., 2017; The Tabula Muris Consortium et al., 2019). In recent years, machine learning methods have been proposed to learn a compact latent representation to address gene expression denoising and data integration (Lopez et al., 2018; Eraslan et al., 2019), and perturbation modeling (Lotfollahi et al., 2019b;a). However, current methods are not able to incorporate prior-knowledge into their learning algorithms. This work focuses on representation learning by exploiting prior-knowledge for single-cell data. Differences in gene expression between cells can be decomposed into observed and unobserved factors. These factors can include non-biological factors such as batch effect, or biological factors, which can often be related to existing knowledge about biological processes and pathways. Recently, (Svensson et al., 2020) introduced an interpretable factor model in the form of a linear autoencoder, however, these authors do neither include prior knowledge, nor aim at explaining variation using prior knowledge.

Among existing models, only the factorial single-cell latent variable model (Buettner et al., 2017) can jointly infer factors that capture different sources of single-cell transcriptome variations, including i) variation in expression attributable to pre-annotated sparse gene sets representing biological knowledge, ii) effects due to additional sparse factors that are meant to explain biological effects, and iii) dense factors that are expected to affect the expression profile of the majority of genes, and often represent technical confounders. This model also allows the assignment of genes to each annotated factor to be refined in a data-driven manner. By that, it offers a powerful alternative to traditional enrichment techniques, which are used to contextualize differential expression signatures with biological knowledge.

In f-scLVM, deterministic approximate Bayesian inference based on variational methods is used to approximate the posterior over all random variables of the model. The f-scLVM python package (slalom) is a custom implementation of the variational Bayesian scheme developed for the model, and we use it as main reference for our benchmarks. It is important to note that the variational Bayesian formulation of the method affects its scalability, and complicates the inference of values of the latent factors for new and out-of-sample data points after the model was already trained.

Here, we present a scalable alternative to f-scLVM to learn latent representations of single-cell RNA-seq data that exploit prior knowledge such as Gene Ontology, resulting in interpretable factors. Our frequentist alternative also allows easier inference of the latent factors for out-of-sample data.

## 2. Model

Our autoencoder can be formulated as:

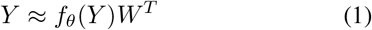

Where *f_θ_*(*Y*) is a neural network with learnable parameters *θ*. This network takes *Y*, the gene expression matrix of size *N × G*, as an input and produces a matrix of factor loadings with size *N* × *K*. *W* in the formula above is a learnable parameter matrix of size *G × K*. A column of *W* corresponds to weights for all genes for a given factor. Analogously, a row of *f_θ_*(*Y*) corresponds to loadings of all factors for a given cell. Also, note that the model has the nonlinear encoder *f_θ_*(*Y*) and the linear decoder *zW^T^*.

After the training phase, the encoder *f_θ_*(*Y*) can also be used to project out-of-sample data to the learned latent space.

If there are observed covariates *X*, i.e. features that are known prior to training, such as batch or cell type. They can be incorporated additively into the model 1 as

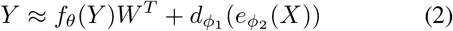

Where *e*_*ϕ*_2__(*X*) is an encoder network taking covariates matrix *X* and *d*_*ϕ*_1__(·) is a decoder network. In particular the linear model can be used in place of the decoder and encoder for the covariates *X*, that is 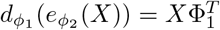.

Only the form 1 (the model without observed covariates) will be discussed further.

There are three kinds of factors in the model.

1. **Annotated factors** correspond to gene sets from pathway databases, such as MSigDB (Subramanian, 2005) or Reactome (Jassal et al., 2019). For these factors we enforce structured sparsity informed by the gene sets.
2. **Sparse unannotated factors** represent biologically meaningful factors that don’t have annotations. These factors are assumed to be generically sparse.
3. **Dense factors** correspond to effects on the expression of large numbers of genes, no sparsity is enforced for this type.

To model annotated and unannotated factors in *W*, we need to introduce the right kind of structured sparsity regularization for the columns of *W*.

The decomposition 1 is fitted to *Y* using regularized L2 loss

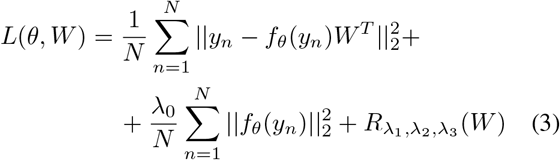

Where λ_0−3_ are regularization hyperparameters, the term *R*_λ_1_,λ_2_,λ_3__(*W*) is a sparsity inducing regularization function, and *y_n_* denotes the n-th row of the data matrix *Y* (*n*-th cell). *L*(*θ,W*) can be optimized using the mini-batch stochastic gradient descent algorithm.

*R*_λ_1_,λ_2_,λ_3__(*W*) in 3 consists of two additive terms

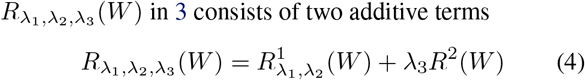

The first term 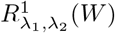 induces structured sparsity on the level of individual genes in each factor (individual elements in each column of *W*).

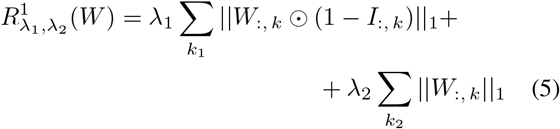

Where *k*_1_, *k*_2_ correspond to annotated factors and sparse unannotated factors respectively. W_:, *k*_ ʘ (1 − *I*_:, *k*_) means Hadamard (element-wise) product between the k-th column of the factor weights matrix W and the k-th column of the binary annotation matrix *I*. The matrix *I*, which is used for the annotated factors, can be formed from pathway databases, such as Reactome or MSigDB, with *I_g,k_* = 1 if the gene *g* is present in the pathway k and *I_g,k_* = 0 otherwise. This means that ||W_:, *k*_ ʘ (1 − *I*_:, *k*_)||_1_ equals to the sum of the absolute values of weights for genes in the factor *k* that are inactive in the annotation of the factor *k* (genes that have *I_g,k_* = 0).

The second regularization term *R*^2^(*W*) in 4 is responsible for deactivating unneeded factors. We use group lasso for this.

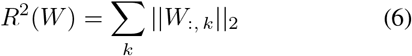

The group lasso ensures that all genes in a factor (a column of the matrix W) are either included or excluded from the model together. This penalty exploits the nondifferentiability of ||*W*_:, *k*_||_2_ at *W*_:, *k*_ = 0; setting whole columns of W to exactly 0.

To minimize 3, the stochastic proximal gradient algorithm is used. This algorithm is employed here because the ordinary stochastic gradient descent algorithm doesn’t account for the points of non-differentiability in *R*_λ_1_,λ_2_,λ_3__(*W*); thus, it can’t enforce the required structured sparsity.

Lets denote the part of the objective function *L*(*θ, W*) (3) without the regularization *R*_λ_1_,λ_2_,λ_3__(*W*) by *F*(*θ, W*). Then, to minimize the objective function 3, we use the update scheme

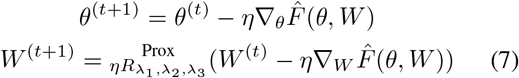

Where 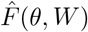 denote the function *F*(*θ, W*) calculated for a mini-batch of samples, *η* is a learning rate. 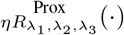 denotes the proximal operator of *R*_λ_1_,λ_2_,λ_3__(*W*)

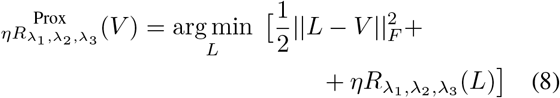

Using the result from (Yu, 2013), it can be shown that the proximal operator 8 has a closed-form expression, which is a composition of the closed-form expressions for the proximal operators corresponding to 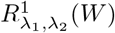 and λ_3_*R*^2^(*W*);.

## 3. Results

To validate the model, we considered a dataset where some sources of variation are known. In (Kang et al., 2017) 8 Lupus patients were stimulated with interferon (IFN) *β*. We expect to see upregulation of the pathways related to interferon signaling.

The autoencoder with only annotated factors was trained on the dataset from the paper with 1k highly variable genes selected, using the Reactome database for the annotated factors. The scatter plots of factor loadings of the annotated factors corresponding to the gene sets from Reactome “INTERFERON SIGNALLING” and “SIGNALING BY THE B CELL RECEPTOR BCR” (Figure 1a) show clear separation of stimulated cells from control cells.

Next, we sought to analyze celltype-specific pathways by using the pathway related to B cells. We can also see a clear separation of B cells from the rest (Figure 1b).

**Figure 1.**
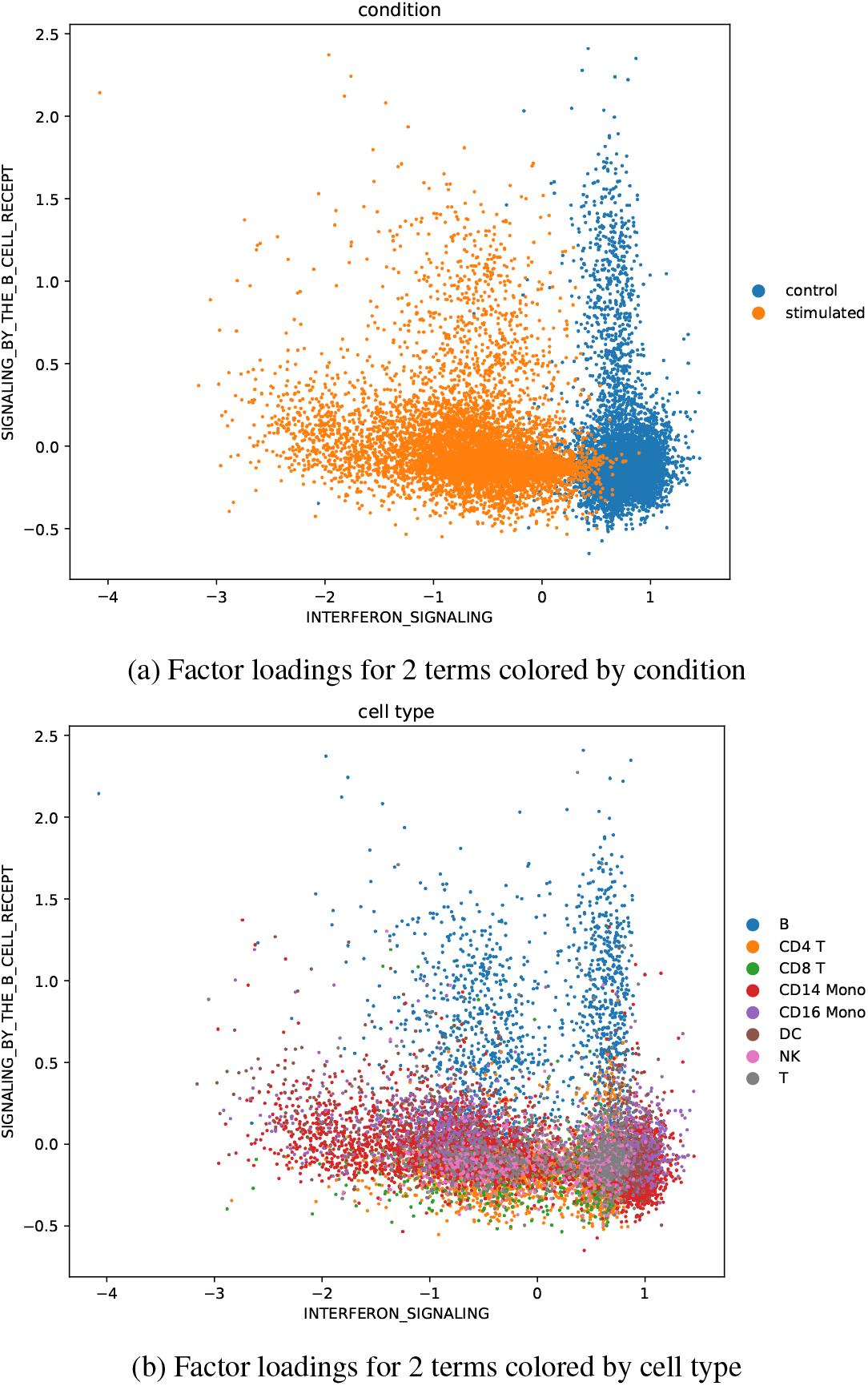
Factor loadings in the interpretable AE.

Also we compared our results with f-scLVM. The python implementation of f-scLVM (slalom) was used for the comparison. In the table below the time to train both models is given.

**Table 1.** Runtime comparison in minutes.

|  | Kang17<br>13.5k cells | Macosko15<br>13.5k cells |
| --- | --- | --- |
| f-scLVM (slalom) | 2203 | 1629 |
| Interpretable AE (this work) | 8 | 7 |

The time is provided for the 13.5k cell subset of the retina dataset from Macosko et al. (2015) and for the dataset from Kang et al. (2017). For both datasets the genes that are not in the annotations from Reactome were filtered out, and also highly variable genes were selected to limit the number of genes to about 1k. It can be seen from the table that slalom requires 270 times as long as the interpretable AE. More importantly, while it is still manageable for datasets of this size, it becomes near impossible to use slalom for much larger datasets, which are common in single cell genomics.

In order to check how well both models explain variation in the Kang et al. (2017) dataset, several terms related to interferon beta were selected, and their loadings were used individually to train uni-variate binary logistic regression on the experimental condition (stimulated vs. control) as class label (Figure 2). Two setups of f-scLVM were used for comparison: f-scLVM without any dense and sparse unannotated factors and f-scLVM with 3 dense factors (default setting). The autoencoder provides better accuracy, and thus better separation, across all factors except for “ANTIVIRAL_MECHANISM_BY_IPN_STIMULATED_GENES”.

**Figure 2.**
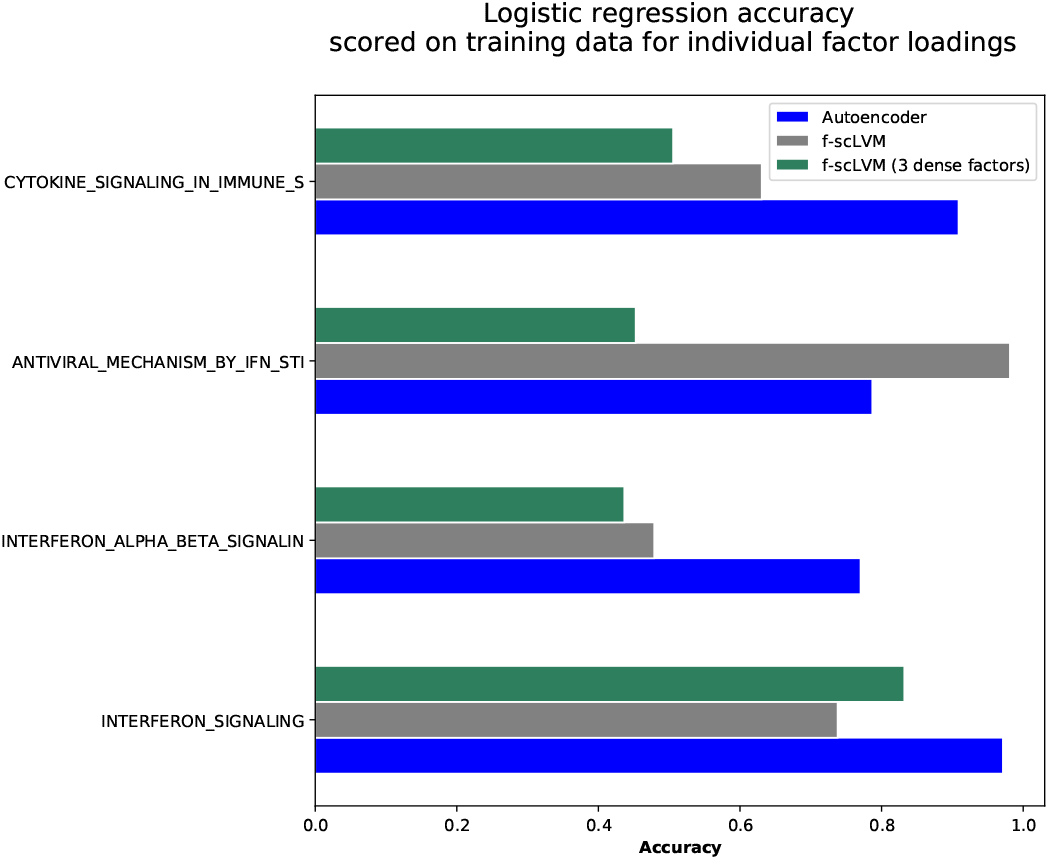
Accuracies of logistic regressions trained on loadings of individual factors vs condition (control, stimulated) for 3 models - autoencoder, f-scLVM without dense and sparse unannotated factors, f-scLVM with 3 dense factors (default setting).

The figure 2 shows the resulting accuracies for training data of logistic regressions trained with the loadings of the individual factors. Two setups of f-scLVM were used for comparison - f-scLVM without any dense and sparse unannotated factors and f-scLVM with 3 dense factors (standard settings). The autoencoder provides better accuracy, and thus better separation, across all factors except for “ANTIVIRAL MECHANISM BY IFN STIMULATED GENES”.

It is interesting that for both autoencoder model and f-scLVM with three dense factors the term which gives the highest accuracy (the best separating term) is “INTERFERON SIGNALLING” (see Figure 2 for the accuracies of classification, Figure 1a for the visualization). However, f-scLVM without dense factors selects “ANTIVIRAL MECHANISM BY IFN STIMULATED GENES” for the separation (Figure 2). It is not clear why training f-scLVM with dense factors, which should account for confounding sources of variation, leads to significant changes in the loadings of the factors which should meaningfully explain biological variation in the data.

Next, we applied our model on a recent, comprising of immune and epithelial cells collected via bronchoalveolar lavage from healthy controls, and patients with moderate and severe COVID-19 Liao et al. (2020). The 10 most important pathways, explaining the variation in the dataset, by L2 norm of their parameter vectors (columns of W in 1) are

1. IMMUNE_SYSTEM
2. METABOLISM_OF_PROTEINS
3. HEMOSTASIS
4. TCA_CYCLE_AND_RESPIRATORY_ELECTRON_TRANSPORT
5. ADAPTIVE_IMMUNE_SYSTEM
6. 3_UTR_MEDIATED_TRANSLATIONAL_REGULATION
7. TRANSLATION
8. METABOLISM_OF_LIPIDS_AND_LIPOPROTEINS
9. INTERFERON_SIGNALING
10. MRNA_PROCESSING

In order to validate interpretability of the learned latent space of pathways, we again investigated factor loadings of the “INTERFERON SIGNALING” term.

The figure 3 shows the separation of control and activated immune cells in moderate and sever COVID-19 in “INTERFERON_SIGNALING” term.

**Figure 3.**
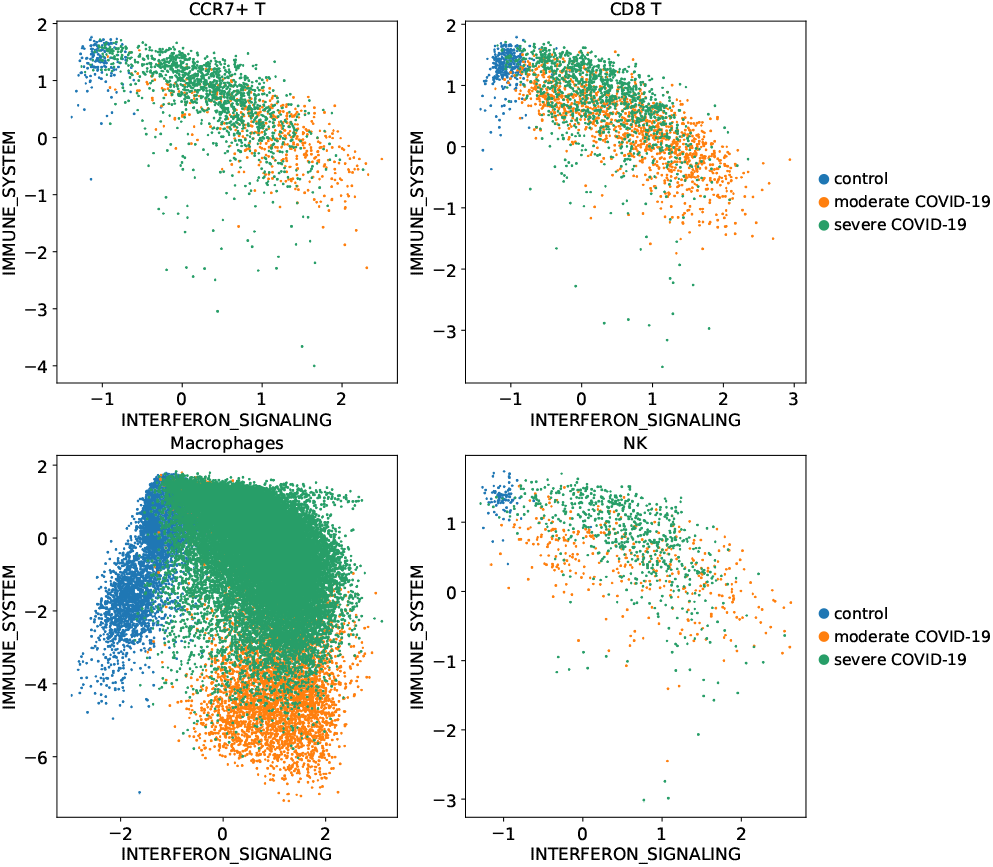
Factor loadings of 2 terms for the four cell types from the COVID-19 dataset

## 4. Discussion and future work

We presented an end-to-end trained autoencoder that can exploit biological knowledge from various databases (Subramanian, 2005; Jassal et al., 2019) to learning interpretable representations of data.

The current model can be extended by learning a more powerful decoder. This requires adding non-linearity to the decoder to learn more complex representations. However, non-linear functions can not easily be incorporated into to model while preserving feature factorization. Gradientbased feature importance methods have been shown to be applicable in this context for image data (Erion et al., 2019). We leave further extensions in this direction for future work.

The code for the model and the accompanying data can be obtained from https://github.com/theislab/InterpretableAutoencoders.

## Notes

### Competing Interest Statement

The authors have declared no competing interest.

## References

The factorial single-cell latent variable model (slalom). https://github.com/bioFAM/slalom.

Buettner, F., Pratanwanich, N., McCarthy, D., Marioni, J., and Stegle, O. f-sclvm: Scalable and versatile factor analysis for single-cell rna-seq. Genome Biology, 18, 12 2017. doi: 10.1186/s13059-017-1334-8.

Eraslan, G., Simon, L. M., Mircea, M., Mueller, N. S., and Theis, F. J. Single-cell RNA-seq denoising using a deep count autoencoder. Nature Communications, 10 (1):390, January 2019. ISSN 2041-1723. doi: 10.1038/s41467-018-07931-2. URL https://www.nature.com/articles/s41467-018-07931-2. Number: 1 Publisher: Nature Publishing Group.

Erion, G. G., Janizek, J. D., Sturmfels, P., Lundberg, S. M., and Lee, S.-I. Learning explainable models using attribution priors. ArXiv, abs/1906.10670, 2019.

Jassal, B., Matthews, L., Viteri, G., Gong, C., Lorente, P., Fabregat, A., Sidiropoulos, K., Cook, J., Gillespie, M., Haw, R., Loney, F., May, B., Milacic, M., Rothfels, K., Sevilla, C., Shamovsky, V., Shorser, S., Varusai, T., Weiser, J., and D’Eustachio, P. The reactome pathway knowledgebase. Nucleic acids research, 48, 11 2019. doi: 10.1093/nar/gkz1031.

Kang, H. M., Subramaniam, M., Targ, S., Nguyen, M., Maliskova, L., McCarthy, E. A., Wan, E., Wong, S. L., Byrnes, L. E., Lanata, C. M., Gate, R. E., Mostafavi, S., Marson, A., Zaitlen, N., Criswell, L. A., and Ye, C. J. Multiplexed droplet single-cell rna-sequencing using natural genetic variation. In Nature Biotechnology, 2017.

Liao, M., Liu, Y., Yuan, J., Wen, Y., Xu, G., Zhao, J., Cheng, L., Li, J., Wang, X., Wang, F., Liu, L., Amit, I., Zhang, S., and Zhang, Z. Single-cell landscape of bronchoalveolar immune cells in patients with covid-19. Nature Medicine, 26, 06 2020. doi: 10.1038/s41591-020-0901-9.

Lopez, R., Regier, J., Cole, M., Jordan, M., and Yosef, N. Bayesian Inference for a Generative Model of Transcriptome Profiles from Single-cell RNA Sequencing. preprint, Bioinformatics, March 2018. URL http://biorxiv.org/lookup/doi/10.1101/292037.

Lotfollahi, M., Naghipourfar, M., Theis, F. J., and Wolf, F. A. Conditional out-of-sample generation for unpaired data using trVAE. arXiv:1910.01791 [cs, eess, q-bio, stat], October 2019a. URL http://arxiv.org/abs/1910.01791. arXiv: 1910.01791.

Lotfollahi, M., Wolf, F. A., and Theis, F. J. scGen predicts single-cell perturbation responses. Nature Methods, 16(8):715–721, August 2019b. ISSN 1548-7091, 1548-7105. doi: 10.1038/s41592-019-0494-8. URL http://www.nature.com/articles/s41592-019-0494-8. tex.ids: lotfollahiScGen-PredictsSinglecell2019a Number: 8 Publisher: Nature Publishing Group.

Macosko, E., Basu, A., Satija, R., Nemesh, J., Shekhar, K., Goldman, M., Tirosh, I., Bialas, A., Kamitaki, N., Martersteck, E., Trombetta, J., Weitz, D., Sanes, J., Shalek, A., Regev, A., and Mccarroll, S. Highly parallel genome-wide expression profiling of individual cells using nanoliter droplets. Cell, 161:1202–1214, 05 2015. doi: 10.1016/j.cell.2015.05.002.

Regev, A., Teichmann, S. A., Lander, E. S., Amit, I., Benoist, C., Birney, E., Bodenmiller, B., Campbell, P., Carninci, P., Clatworthy, M., et al. Science forum: the human cell atlas. Elife, 6:e27041, 2017.

Subramanian, A. Gene set enrichment analysis: a knowledge-based approach for interpreting genome-wide expression profiles. proc natl acad sci u s a. Proceedings of the National Academy of Sciences, 102:15545–15550, 10 2005.

Svensson, V., Gayoso, A., Yosef, N., and Pachter, L. Interpretable factor models of single-cell rna-seq via variational autoencoders. Bioinformatics (Oxford, England), 03 2020. doi: 10.1093/bioinformatics/btaa169.

The Tabula Muris Consortium, Pisco, A. O., McGeever, A., Schaum, N., Karkanias, J., Neff, N. F., Darmanis, S., Wyss-Coray, T., and Quake, S. R. A Single Cell Transcriptomic Atlas Characterizes Aging Tissues in the Mouse. preprint, Cell Biology, June 2019. URL http://biorxiv.org/lookup/doi/10.1101/661728.

Yu, Y. On decomposing the proximal map. In Proceedings of the 26th International Conference on Neural Information Processing Systems - Volume 1, NIPS’13, pp. 91–99, Red Hook, NY, USA, 2013. Curran Associates Inc.

